# A Single Cell Gene Expression Atlas of 28 Human Livers

**DOI:** 10.1101/2020.12.23.422515

**Authors:** Joseph Brancale, Sílvia Vilarinho

## Abstract

The liver is the largest solid organ in the human body and is responsible for a multitude of essential functions for survival. Chronic liver injury affects over 1 billion people worldwide and therapeutic options other than liver transplantation are a critical unmet medical need. Thus, advances in molecular hepatology are essential to facilitate the discovery of new therapeutic targets. Here we describe the aggregation and integration of single cell RNA-sequencing in more than 36,000 cells from 28 human livers reported in five independent studies. Noteworthy, the merged data shows a high degree of overlap, demonstrating the robustness of liver gene expression at single cell level independent of age, gender, liver collection, processing and sequencing methods. Hence, this data allowed us to develop a user-friendly web browser for quick and easy interrogation and comparison of gene expression across a variety of parenchymal and non-parenchymal liver cell populations. Collectively, this study provides the largest human liver transcriptomic single cell atlas accessible for interactive visualization via an open-access web portal to the research community worldwide.

## INTRODUCTION

Chronic liver disease (CLD) affects over 1 billion people worldwide(1) and therapeutic options other than liver transplantation are an unmet medical need. Thus, advances in molecular hepatology are essential to facilitate the discovery of new therapeutic targets. Next generation sequencing technologies have revolutionized our understanding of human health and disease through its application to DNA and RNA sequencing. The latter comprises single-cell transcriptomics, which enables to sequence thousands of cells in a single sample and to analyze the gene expression of all cells individually. This creates an unbiased approach to assess cell identity and heterogeneity and to uncover rare cell populations otherwise obscured in bulk RNA-sequencing studies. Taken advantage of fully commercialized workflows, high-throughput single-cell RNA-sequencing (scRNA-seq) has been increasingly used to investigate the cell complexity and heterogeneity of human organs, including the liver. Whereas access to good quality fresh human healthy liver tissue has represented a major limitation in the field, in the last two years, five independent studies have analyzed human healthy livers at single cell resolution(2–6). Thus, integration of these five scRNA-seq datasets can provide further insight into the transcriptomic architecture and stability of the human liver in physiological conditions, representing highly valuable, and previously unattainable, information to the liver research community worldwide. Here, we successfully integrate and analyze available human liver scRNA-seq data and provide an interactive open-access online cell browser platform for easy access of gene expression across a variety of annotated parenchymal and non-parenchymal cells derived from 28 human healthy livers.

## METHODS

Data was accessed at the NCBI GEO using the accession numbers GSE115469, GSE136103, GSE129933, GSE124395, GSE130473. For GSE136103, GSE129933, GSE124395, GSE130473, the data was downloaded from GEO and imported directly into Seurat v.3.2.2 (7) using the barcode, gene expression, and annotation files. For GSE115469, raw sequencing files were downloaded and processed through CellRanger (v. 3.0.2) using default settings before imported into Seurat. Once imported into Seurat, each dataset was individually filtered (transcript present in 3 cells minimum, between 250-1000 transcripts, and mitochondrial sequences < 30%) and normalized. For each dataset 2,000 variable features were identified using the variance stabilizing transformation (VST) method(8). In order for Seurat integration to be performed a k filter of 94 was applied (maximum acceptable filter) and anchor genes were identified and used for integration with a dimensionality of 30. The data was scaled and PCA and UMAP were run for dimensionality reduction using the same variable of 30. Cluster identification was performed using a shared nearest neighbors-based algorithm with a resolution of 0.5. Marker genes were identified and used to identify cluster cell types. An R Shiny application was created for interactive visualization and can be accessed at http://liveratlas-vilarinholab.med.yale.edu/.

## RESULTS

We report the integration of all available scRNA-seq datasets (Figure 1A-C) derived from human healthy livers into the largest human liver single cell atlas. This cumulative dataset encompasses 28 livers from five independent studies (Table 1). Liver samples were obtained from individuals of both genders, a wide range of age groups (21 to 65 years of age) and with a variety of underlying medical conditions (Table 1). Despite the potential technical effects associated with distinct cell collection and processing methods, batches and sequencing platforms, we found a high level of consistency in clustering of different liver cell populations among these 28 livers (Figure 1D).

**Table 1.**
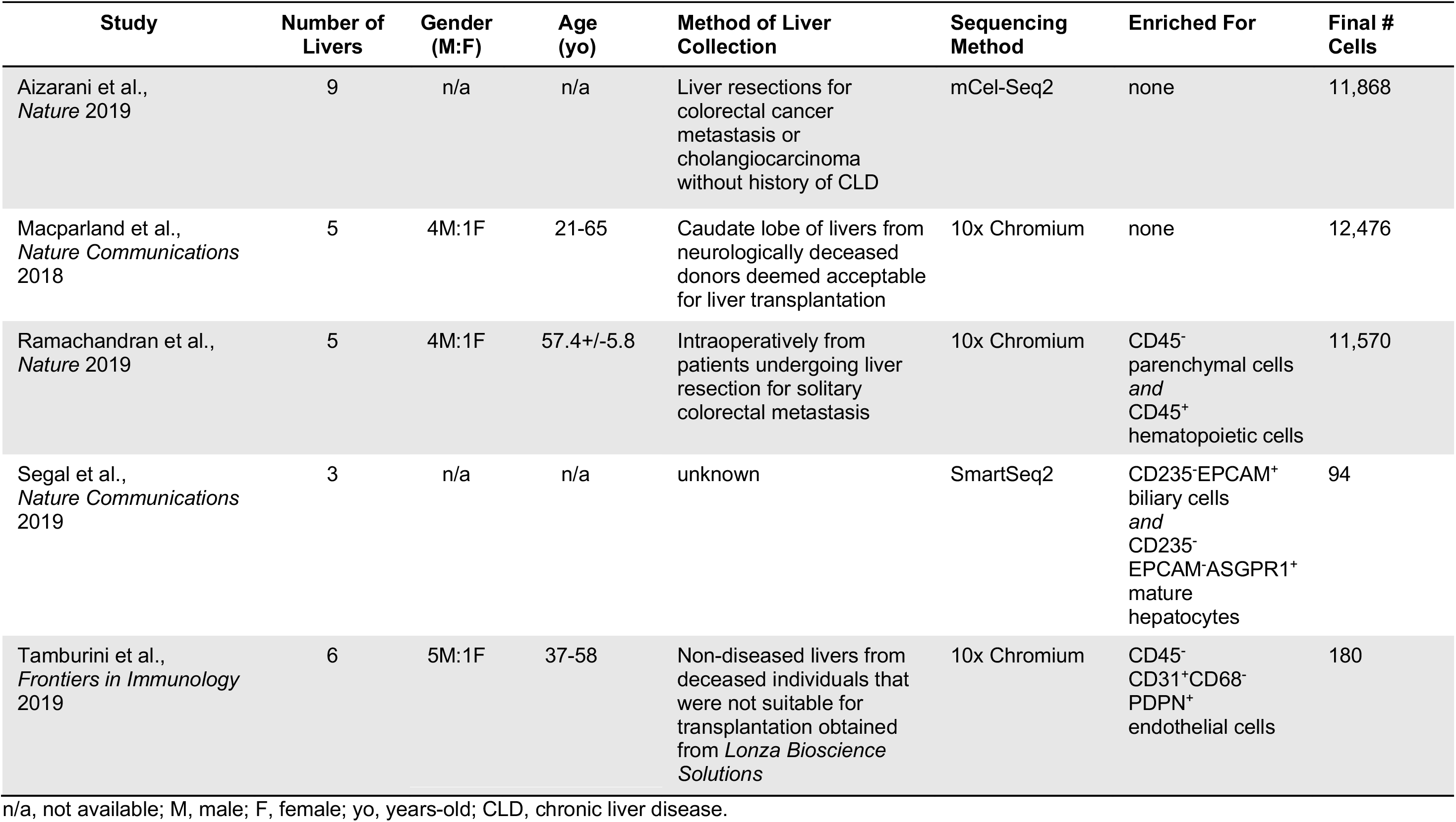
Summary of the demographics, collection and sequencing methods of the healthy human liver samples reported in five independent studies.

**Figure 1.**
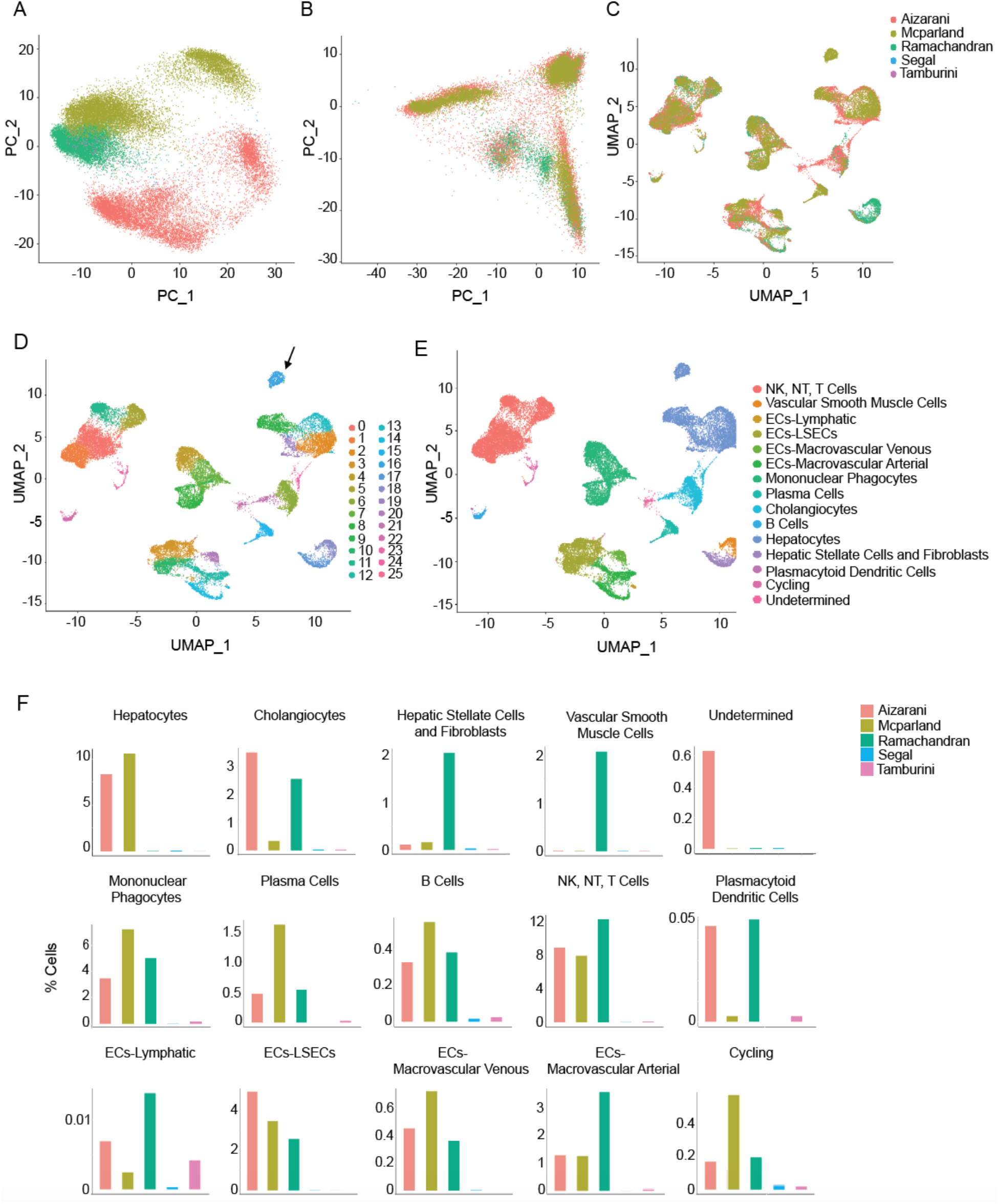
Aggregation and integration of five non-diseased human liver single cell RNA-sequencing datasets. (A) Principle component analysis of individual datasets before integration. (B) Principle component analysis after integration using Seurat version 3. (C) UMAP plot of integrated cells clustered into cell types among the five different studies. (D) UMAP plot of merged 36,188 cells colored by clusters. Arrow indicates cluster 16, which is nearly exclusively detected in one out of the 28 livers analyzed. (E) UMAP plot of merged 36,188 cells colored by cell types. (F) Proportion of the 36,188 cells across each paper and each cell (sub)type.

In aggregate, our merged dataset comprises 26 clusters of a total of 36,188 liver cells (Figure 1D). Cell lineage was inferred from a combination of unbiased clustering and gene expression profiles. It is composed by 6,895 hepatocytes, 2,357 cholangiocytes, 6,876 liver endothelial cells, 1,604 mesenchymal cells and 18,223 immune cells (Figure 1E,F). There were 234 cells that remain undetermined. A more granular annotation of endothelial cells led to its subclassification into four subsets: (i) liver sinusoidal endothelial cells expressing *DNASE1L3* and *CLEC4G*; (ii) macrovascular-venous endothelial cells expressing *RSPO3*; (iii) macrovascular-arterial endothelial cells expressing *EHD4*; and (iv) lymphatic endothelial cells expressing *CCL21A*, *PRSS23*, *IFI27L2A* and *TIMP2A* (Figure 1E,F). Mesenchymal cells were classified into two subgroups: (i) hepatic stellate cells (HSC)/fibroblasts expressing *RGS5* and (ii) vascular smooth muscle cells expressing *MYH11*.

Based on the methods of cell isolation, two studies were enriched for certain cell populations, such as mesenchymal cells(6) or liver endothelial cells(5), and our merged data detect clustering of these cell populations as expected (Figure 1D-F). This large dataset also allowed to identify cluster 16 as composed almost exclusively from hepatocytes isolated from a single individual reported in the MacParland et al. study (Figure 1D).

Importantly, our efforts to gain further insight into the molecular architecture and robustness of gene expression in the liver by successfully merging over 36,000 good quality cells from 28 healthy human livers led to the generation of an open-access web browser of the largest human liver single cell atlas available at: http://liveratlas-vilarinholab.med.yale.edu/. This new resource facilitates user-friendly access and interactive visualization of gene expression for each cell (sub)type, which encompasses a range of abundant to rare liver cell populations. Moreover, this new web tool also provides information on the proportion of cells where a certain gene of interest is detected amongst the dominant cell type and in other liver cell populations.

## DISCUSSION

The major contribution of this study is the generation of an open access web browser that allows easy access of scRNA-sequencing datasets and interactive visualization of gene expression in 36,188 human liver cells isolated from 28 individuals with no chronic liver disease. This resource tool has the potential to accelerate basic, translational and clinical research in liver health and disease globally since it provides the largest amount of good quality liver gene expression information at single cell level accessible to the research community independent of investigators’ computer science training and skills.

Importantly, the success of this data merge demonstrates the stability of gene expression in the liver at single cell level across individual differences and variations in collection, processing and sequencing methods. This data shows overall little variation in the transcriptomic profile of non-diseased livers, suggesting the liver’s capability to endure perturbation to homeostasis. This raises an interesting possibility that the reduced interpersonal gene expression variability detected in the liver at the single cell level might contribute in part to the fact that liver transplant recipients demonstrate lower incidence of chronic rejection and require lower amount of initial and maintenance of immunosuppression therapy as compared to any other solid organ transplant recipients.

By combining five datasets, this study also allowed to identify a cluster comprised almost exclusively by hepatocytes from a single individual, raising the suspicion that these transcriptional changes might represent an outlier and not a cluster regularly detected in human healthy livers.

Collectively, this study provides a valuable online resource for gene expression search and comparison across the diverse human liver cell populations. The web browser was designed to deliver up-to-date single cell gene expression data accessible to any researcher worldwide working on liver biology research questions independent of their level of bioinformatic skills.

## ACKNOWLEDGMENTS

Our laboratory is supported by the Doris Duke Charitable Foundation Grant #: 2019081 and the National Institute of Diabetes and Digestive and Kidney Diseases of the National Institute of Health under Award Number K08 DK113109. J.B. is supported by the National Institute of General Medical Sciences of the National Institute of Health under Award Number 1T32GM136651-01. The content is solely the responsibility of the authors and does not necessarily represent the official views of the National Institute of Health.

